# Alcohol tolerance encoding in sleep regulatory circadian neurons in Drosophila

**DOI:** 10.1101/2023.01.30.526363

**Authors:** Anthony P. Lange, Fred W. Wolf

## Abstract

Alcohol tolerance is a simple form of behavioral and neural plasticity that occurs with the first drink. Neural plasticity in tolerance is likely a substrate for longer term adaptations that can lead to alcohol use disorder. Drosophila develop tolerance with characteristics similar to vertebrates, and it is useful model for determining the molecular and circuit encoding mechanisms in detail. Rapid tolerance, measured after the first alcohol exposure is completely metabolized, is localized to specific brain regions that are not interconnected in an obvious way. We used a forward neuroanatomical screen to identify three new neural sites for rapid tolerance encoding. One of these was comprised of two groups of neurons, the DN1a and DN1p glutamatergic neurons, that are part of the Drosophila circadian clock. We localized rapid tolerance to the two DN1a neurons that regulate arousal by light at night, temperature-dependent sleep timing, and night-time sleep. Two clock neurons that regulate evening activity, LNd6 and the 5th LNv, are postsynaptic to the DN1as and they promote rapid tolerance via the metabotropic glutamate receptor. Thus, rapid tolerance to alcohol overlaps with sleep regulatory neural circuitry, suggesting a mechanistic link.

## Introduction

Alcohol Use Disorder (AUD) is a progressive, chronic, and recurring brain disease that causes extraordinarily long-term changes to brain function. Multiple forms of behavioral adaptations to ethanol, the active ingredient in alcohol, occur that are mostly defined operationally. Determining their relative importance for AUDs and their interconnectedness will help determine the longitudinal and spatial pathways in addiction. Longer term forms of adaption to ethanol likely build on simpler early forms. Early adaptations are amenable to complete molecular and neural circuit definition.

Ethanol tolerance is a simple and early adaptation that is defined as the acquired resistance to the pharmacological effects of the drug; tolerance can facilitate increased intake. Tolerance is classically divided into three forms: acute (acquired within a drinking session), rapid (expressed after the first drink is completely metabolized), and chronic (Fadda and Rossetti, 1998). The fly Drosophila exhibits all three forms of tolerance (Scholz et al., 2000; Berger et al., 2004). Rapid tolerance is currently the best characterized form in Drosophila (**Figure 1A**). An acute just sedating dose of ethanol results in tolerance to its inebriating and sedating properties, sensitization to its locomotor activating properties, and it primes flies for developing ethanol preference (**Figure 1B**) (Scholz et al., 2000; Kong et al., 2010a; Peru Y Colón de Portugal et al., 2014). Molecular parallels to early forms of ethanol-induced neural plasticity in mammals exist (Cowmeadow et al., 2005; Morozova et al., 2006; Kong et al., 2010a; Sakharkar et al., 2012; Ghezzi et al., 2013b; Engel et al., 2016; Berkel and Pandey, 2017; Ranson et al., 2020).

**Figure 1.**
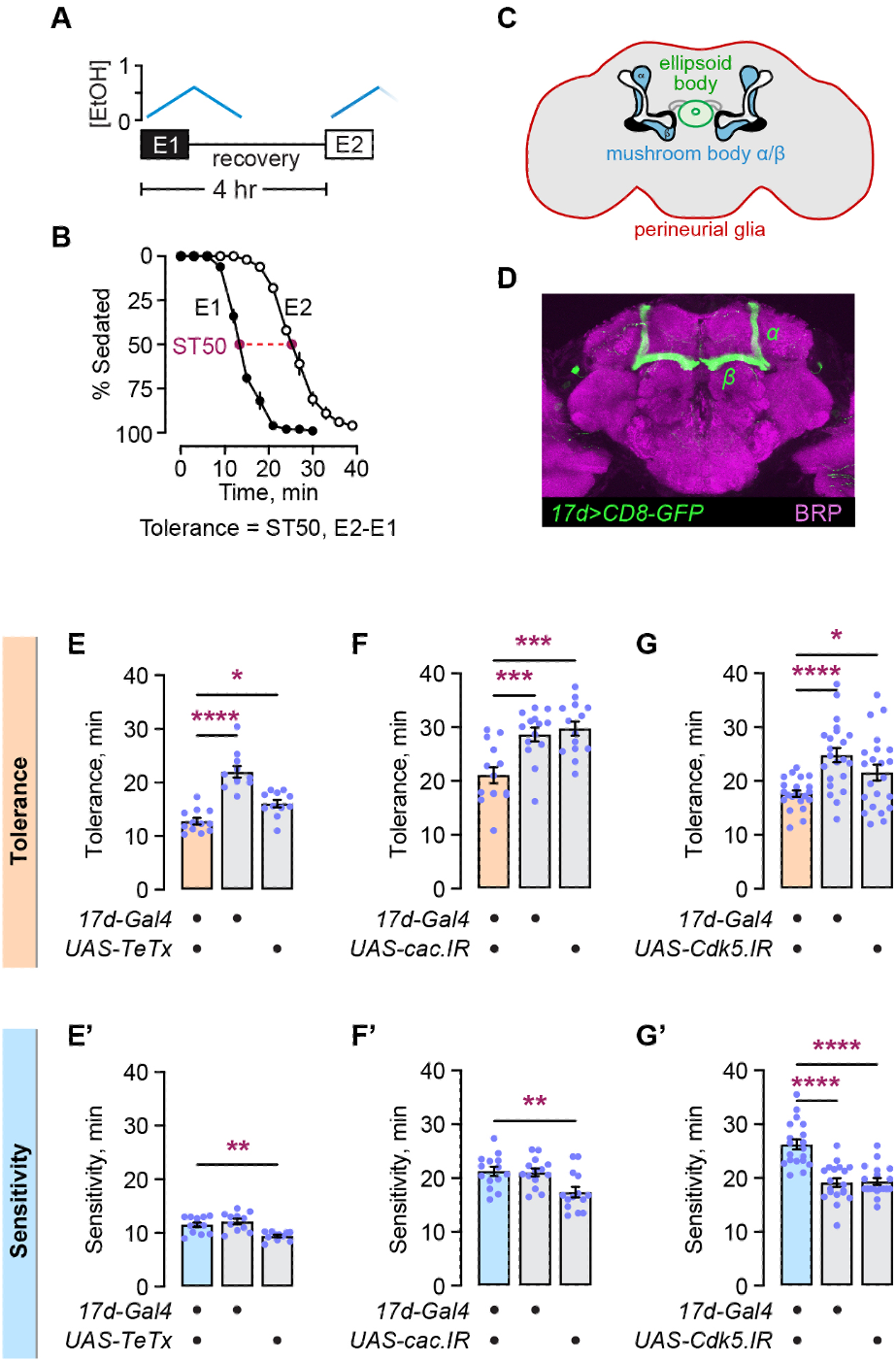
Presynaptic genes promote rapid ethanol tolerance in the mushroom body α/β lobe intrinsic neurons. **A**. Scheme for induction and detection of rapid tolerance. Identical ethanol exposures, E1 and E2, are separated by 4 hr and result in identical accumulation and dissipation kinetics for internal ethanol concentrations. **B**. Time course for ethanol sedation, for E1 and E2, and the calculation of ethanol tolerance. **C**. Diagram of the Drosophila brain, depicting the known sites for rapid tolerance encoding. **D**. Expression pattern in the mushroom body α/β neurons of the *17d-Gal4* transgene, detected by the *UAS-CD8-GFP* reporter transgene, and counterstained for the ELKS/CAST ortholog BRP to reveal the synaptic neuropil. **E,E’**. Tolerance (E) and sensitivity (E’) when presynaptic release is blocked by expression of the tetanus toxin light chain (*UAS-TeTx*) in *17d-Gal4* neurons. **F,F’**. Effect of decreasing expression of the Cav2.1 Ca^2+^ channel *cac* in the *17d-Gal4* pattern. **G,G’**. Effect of decreasing expression of the kinase *Cdk5* in the *17d-Gal4* pattern. *Statistics*: **E**-**G’**: Quantitative data are mean±SEM. **E**. One-way ANOVA (*p*<0.0001) with Dunnett’s multiple comparisons, *Gal4*/*UAS* versus *Gal4*: \*\*\*\**p*<0.0001, *Gal4*/*UAS* versus *UAS*: \**p*=0.0129. **E’**. One-way ANOVA (*p*=0.0002) with Dunnett’s multiple comparisons, *Gal4*/*UAS* versus *Gal4*: *p*=0.4929, *Gal4*/*UAS* versus *UAS*: \*\**p*=0.0025. **F**. One-way ANOVA (*p*<0.0001) with Dunnett’s multiple comparisons, *Gal4*/*UAS* versus *Gal4*: \*\*\**p*=0.0007, *Gal4*/*UAS* versus *UAS*: \*\*\**p*=0.0001. **F’**. One-way ANOVA (*p*=0.0034) with Dunnett’s multiple comparisons, *Gal4*/*UAS* versus *Gal4*: *p*=0.9781, *Gal4*/*UAS* versus *UAS*: \*\**p*=0.0049. **G**. Brown-Forsythe ANOVA (*p*=0.0005) with Dunnett’s T3 multiple comparisons, *Gal4*/*UAS* versus *Gal4*: \*\*\*\**p*<0.0001, *Gal4*/*UAS* versus *UAS*: \**p*=0.0394. **G’**. One-way ANOVA (*p*<0.0001) with Dunnett’s multiple comparisons, *Gal4*/*UAS* versus *Gal4*: \*\*\*\**p*<0.0001, *Gal4*/*UAS* versus *UAS*: \*\*\*\**p*<0.0001.

Defining the neural circuit that encodes rapid ethanol tolerance is essential for understanding the mechanisms and contexts of ethanol action. Current anatomical localization of rapid tolerance includes the α/β lobes of the mushroom bodies, the ellipsoid body, the perineurial glia, and some other less well-defined sites (Ghezzi et al., 2013a; Engel et al., 2016; Park et al., 2017; Ruppert et al., 2017; Parkhurst et al., 2018; Kang et al., 2020) (**Figure 1C**). These anatomical sites provide some insight into the nature of ethanol tolerance, based on their characterization in other behaviors. For example, the mushroom bodies are a major site of learning and memory in Drosophila (Modi et al., 2020). The mushroom bodies play multiple roles in ethanol behaviors including the coding of preference and reward learning, consolidation, and retrieval (Kaun et al., 2011; Xu et al., 2012). The ellipsoid body is a compass for flight and locomotor navigation, and it also functions in sleep and arousal state (Seelig and Jayaraman, 2015; Andreani et al., 2022). The perineurial glia form the interface between the brain and the circulatory system; their role in brain physiology is less well understood. These early attempts at building a tolerance circuit were largely limited to well-studied neuropils in the fly brain, whereas 80-90% of neurons in the fly brain are not in these neuropils.

Advances in genetic targeting of individual neurons in the fly brain, coupled with the advent of complete brain connectomics, promises to make the whole tolerance circuit available for characterization (Luan et al., 2020; Galili et al., 2022). However, there exists no short path connectivity between the brain regions known to function in rapid tolerance, and it is not known how tolerance information flows in and out of the defined neurons. For example, output from the mushroom body intrinsic neurons is via the mushroom body output neurons (MBONs) (Rubin 2014). A subset of MBONs synapse onto fan-shaped body neurons, that are in turn synaptically connected to ellipsoid body neurons (Jayaraman 2021). There are other paths that tolerance information could take between these brain structures, and the direction of information flow is not known. Alternatively, the currently known tolerance brain regions could parallel process separable aspects of tolerance. Finding additional tolerance circuitry can help us build better models of tolerance encoding.

We performed a functional anatomical screen for new rapid tolerance neurons in the Drosophila brain. We characterized one of three new sites, uncovering a role for the glutamatergic DN1a circadian clock neurons. The DN1a neurons regulate the timing and quality of evening sleep in relation to environmental inputs, and they promote rapid ethanol tolerance development. Additional clock neurons that control evening behavior are postsynaptic to the DN1as and are required for rapid ethanol tolerance development. Both the mushroom bodies and the ellipsoid body function in sleep, suggesting relationship between sleep and rapid tolerance.

## Results

Rapid ethanol tolerance (hereafter referred to as tolerance) is thought to involve changes in presynaptic function, based in part on changes in expression of genes encoding the Cav2.1 channel Cacophony (*cac*), the presynaptic kinase Cdk5, and the synaptic vesicle regulator Synapsin (Godenschwege et al., 2004; Ghezzi et al., 2013a; Engel et al., 2016). The mushroom body α/β lobes require *Sirt1* for rapid tolerance, and acute ethanol regulation of presynaptic gene expression is lost in *Sirt1* null mutant flies. Hence, we used the mushroom body α/β lobes to test if decreasing expression of *cac* and *Cdk5* affects tolerance development. We first verified that silencing synaptic output from the α/β lobe neurons decreased tolerance, by expressing the tetanus toxin short chain (*UAS-TeTx*) specifically in the α/β lobes using the *17d-Gal4* transgene driver (**Figure 1D**). Silencing synaptic output decreased tolerance and did not affect naive ethanol sensitivity (**Figure 1E,E’**), as previously reported (Engel et al., 2016). Decreasing expression of either *cac* or *Cdk5* in the α/β lobes also decreased tolerance (**Figure 1F,F’,G,G’**). Thus, presynaptic release from the α/β lobe neurons is important for tolerance development.

To identify additional components of the neural circuitry for tolerance, we reasoned that decreasing presynaptic release in sparse patterns of neurons that may contain tolerance neurons would impact tolerance development. We manually selected *enhancer-Gal4* strains from the FlyLight collection that appeared to express in sparse and distinct patterns in the Drosophila brain (Pfeiffer et al., 2008). To improve our chances of discovering new tolerance circuitry, we selected against patterns with *Gal4* expression in either the mushroom bodies or the ellipsoid body. Decreasing *cac* expression in neurons in 112 different *enhancer-Gal4* patterns resulted in either decreased or increased tolerance (**Figure 2A**). We retested 24 of the top hits that were greater than one standard deviation from the mean tolerance difference score. The retests included the full suite of genetic controls and were tested over 8-10 trials. Three *enhancer-Gal4* strains in retest resulted in robust reduction in tolerance when driving *cac* RNAi: *R82F12, R18H11*, and *R79H04* (**Figure 2B,B’**). Blockade of synaptic release with TeTx also reduced tolerance in all three *enhancer-Gal4* patterns, indicating that neurotransmission is required for rapid tolerance in multiple different neurons in the Drosophila brain (**Figure 2C,C’**).

**Figure 2.**
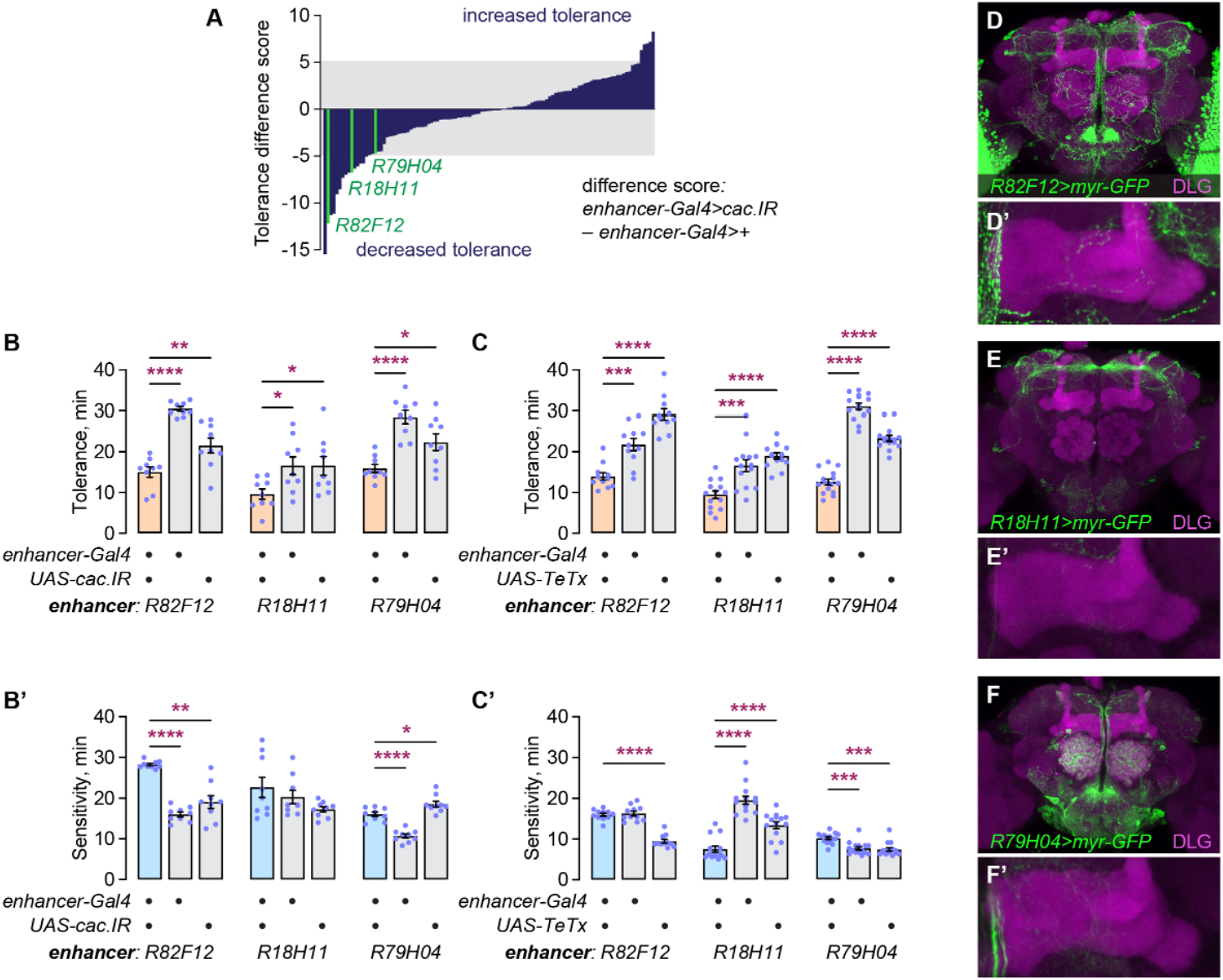
A functional neuroanatomical screen identifies three patterns of neurons that promote rapid tolerance development through presynaptic release. **A**. Tolerance difference score for 112 *enhancer-Gal4* strains expressing RNAi for *cac*. Grey region represents 2 standard deviations from the mean of the difference scores. Highlighted in green are three strains that passed secondary screens and that were further characterized. **B,B’**. Reduction of *cac* expression in *R82F12-Gal4* (left), *R18H11-Gal4* (middle), and *R79H04-Gal4* (right) effects on rapid tolerance (B) and sensitivity (B’). **C,C’**. Effect of tetanus toxin blockade of presynaptic release in the same three *enhancer-Gal4* strains for rapid tolerance (C) and sensitivity (C’). **D**-**F**. Expression pattern of the *enhancer-Gal4* strains, revealed with the *UAS-myr-GFP* plasma membrane-tethered GFP, and counterstained with an antibody to the Discs Large (DLG) synaptic protein. **D’**-**F’**. Enlargement of a substack from D-F encompassing the horizontal lobes of the mushroom bodies. *Statistics*: **B**-**C**: Quantitative data are mean±SEM. **B**. Left panel: One-way ANOVA (*p*<0.0001) with Dunnett’s multiple comparisons, *Gal4*/*UAS* versus *Gal4*: \*\*\*\**p*<0.0001, *Gal4*/*UAS* versus *UAS*: \*\**p*=0.0040. Middle panel: One-way ANOVA (*p*=0.0293) with Dunnett’s multiple comparisons, *Gal4*/*UAS* versus *Gal4*: \**p*=0.0376, *Gal4*/*UAS* versus *UAS*: \**p*=0.0379. Right panel: One-way ANOVA (*p*<0.0001) with Dunnett’s multiple comparisons, *Gal4*/*UAS* versus *Gal4*: \*\*\*\**p*<0.0001, *Gal4*/*UAS* versus *UAS*: \**p*=0.0162. **B’**. Left panel: Kruskal-Wallis test (p=0.0001) with Dunn’s multiple comparisons, *Gal4*/*UAS* versus *Gal4*: \*\*\*\**p*<0.0001, *Gal4*/*UAS* versus *UAS*: \*\**p*=0.0059. Middle panel: One-way ANOVA (*p*=1120). Right panel: One-way ANOVA (*p*<0.0001) with Dunnett’s multiple comparisons, *Gal4*/*UAS* versus *Gal4*: \*\*\*\**p*<0.0001, *Gal4*/*UAS* versus *UAS*: \**p*=0.0126. **C**. Left panel: One-way ANOVA (*p*<0.0001) with Dunnett’s multiple comparisons, *Gal4*/*UAS* versus *Gal4*: \*\*\**p*=0.0003, *Gal4*/*UAS* versus *UAS*: \*\*\*\**p*<0.0001. Middle panel: One-way ANOVA (*p*<0.0001) with Dunnett’s multiple comparisons, *Gal4*/*UAS* versus *Gal4*: \*\*\**p*=0.0001, *Gal4*/*UAS* versus *UAS*: \*\*\*\**p*<0.0001. Right panel: One-way ANOVA (*p*<0.0001) with Dunnett’s multiple comparisons, *Gal4*/*UAS* versus *Gal4*: \*\*\*\**p*<0.0001, *Gal4*/*UAS* versus *UAS*: \*\*\*\**p*<0.0001. **C’**. Left panel: One-way ANOVA (*p*<0.0001) with Dunnett’s multiple comparisons, *Gal4*/*UAS* versus *Gal4*: *p*=0.8707, *Gal4*/*UAS* versus *UAS*: \*\*\*\**p*<0.0001. Middle panel: One-way ANOVA (*p*<0.0001) with Dunnett’s multiple comparisons, *Gal4*/*UAS* versus *Gal4*: \*\*\*\**p*<0.0001, *Gal4*/*UAS* versus *UAS*: \*\*\*\**p*<0.0001. Right panel: Kruskal-Wallis test (*p*<0.0001) with Dunn’s multiple comparisons, *Gal4*/*UAS* versus *Gal4*: \*\*\**p*=0.0007, *Gal4*/*UAS* versus *UAS*: \*\*\**p*=0.0001.

We visualized the neurons in the three *enhancer-Gal4* patterns by expressing plasma membrane-bound GFP (*UAS-myr-GFP*) in each pattern and performing immunohistochemistry and confocal microscopy (**Figure 2D-F**). All three *enhancer-Gal4* patterns expressed in sparse patterns of neurons in the brain, with *R18H11* appearing the sparsest. None expressed in the mushroom bodies, the ellipsoid bodies, or the perineurial glia. *R18H11* contains a subset of the neurons that comprise the circadian circuitry, the DN1 neurons that have cell bodies in the dorsal region of the brain, and this *Gal4* transgene was used previously to characterize the function of these clock neurons (**Figure 2E**) (Kunst et al., 2014). The DN1 neurons form two clusters consisting of two DN1a neurons and seven DN1p neurons (Shafer et al., 2006, 2022).

Neurons in both the DN1a and DN1p clusters are glutamatergic. To ask if glutamatergic neurons are responsible for promoting tolerance, we expressed an RNAi for the glutamate transporter vGlut in each of the three *enhancer-Gal4* patterns. *vGlut* RNAi in *R82F12* and *R18H11* reduced tolerance, whereas it did not in *R79H04* (**Figure 3A, A’**). A second RNAi directed against *vGlut* also reduced tolerance when expressed in *R82F12* neurons, indicating that the effect on tolerance is due to the reduction in vGlut expression (**Figure 3B,B’**). Hence, *R79H04* contains non-glutamatergic tolerance neurons that are distinct from those found in other known tolerance *enhancer-Gal4* patterns. Additionally, it is likely that the glutamatergic DN1 neurons are responsible for promoting ethanol tolerance. To ask if the role of glutamatergic transmission in tolerance is an adult role, we conditionally expressed *vGlut* RNAi in all adult neurons. *Gal80ts* encodes a temperature-sensitive repressor of GAL4, such that it represses GAL4 activity at 18°C and is inactive at 29°C (McGuire et al., 2003). Tolerance was reduced when GAL4 was allowed to be active and drive *vGlut* RNAi only in adult neurons (**Figure 3C,C’**).

**Figure 3.**
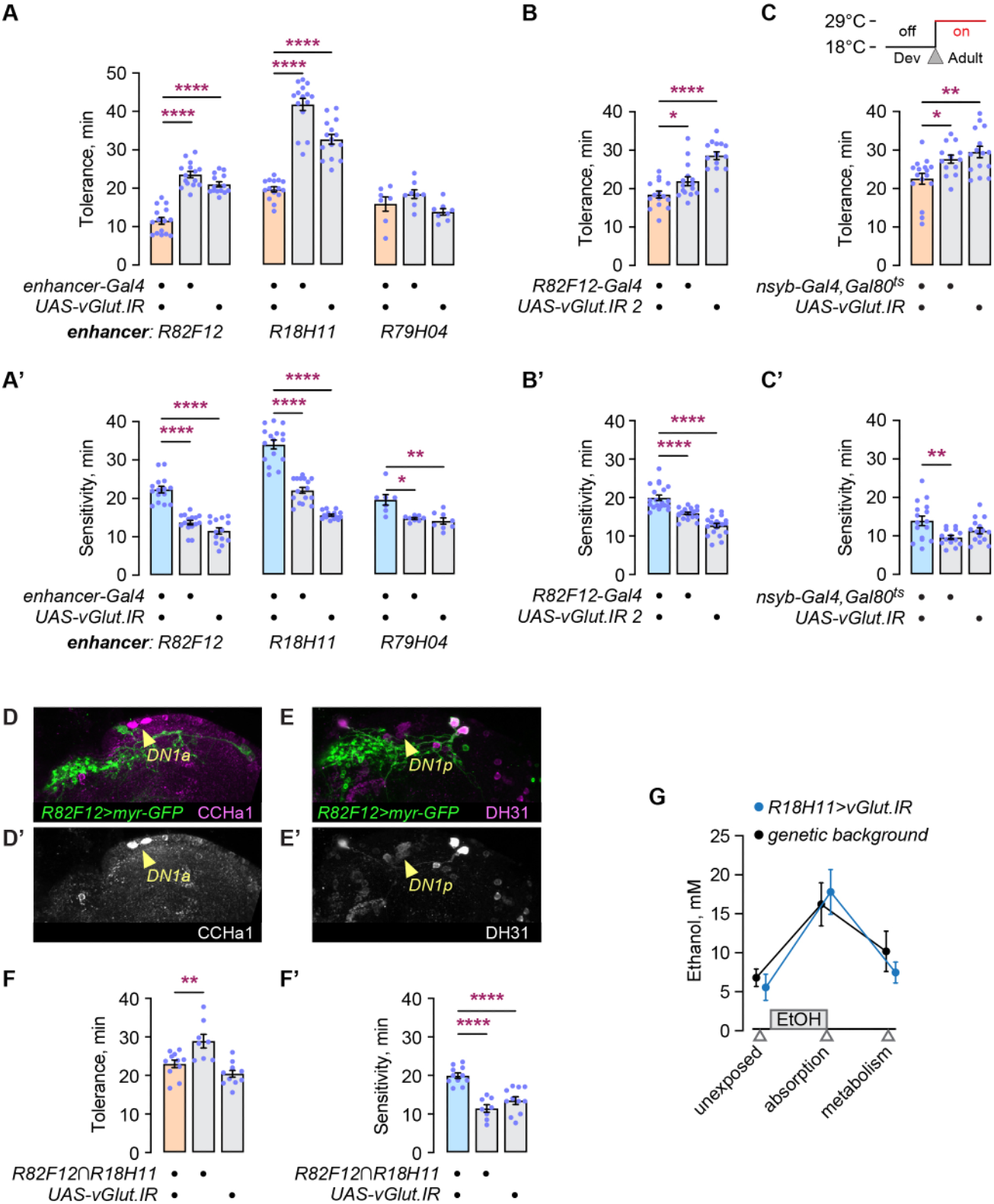
Distinct rapid tolerance neurons exist in each *enhancer-Gal4*, including glutamatergic dorsal clock neurons in *R18H11-Gal4*. **A,A’**. Reduction of glutamate release with vesicular glutamate transporter *vGlut* RNAi effects on rapid tolerance (A) and sensitivity (A’). **B,B’**. Reduction of *vGlut* expression using a second independent RNAi transgene, effects on tolerance (B) and sensitivity (B’) in *R82F12-Gal4* expressing neurons. **C,C’**. Adult specific *vGlut* RNAi in *R82F12-Gal4* expressing neurons, effect on rapid tolerance (C) and sensitivity (C’). **D,D’**. Counterstaining of *R82F12>myr-GFP* brains with anti-CCHa1 to detect DN1a neurons. **E,E’**. Counterstaining of *R82F12>myr-GFP* brains with anti-DH31 to detect DN1p neurons. Micrographs in D and E depict the dorsal hemisphere of an adult brain. **F,F’**. *Split-Gal4* driving expression of *vGlut* RNAi in neurons that are common between *R82F12* and *R18H11*, effect on rapid tolerance (F) and sensitivity (F’). **G**. Effect of *vGlut* RNAi in *R18H11-Gal4* neurons on ethanol absorption and metabolism. *Statistics*: **A**-**C,F**: Quantitative data are mean±SEM. **A**. Left panel: One-way ANOVA (*p*<0.0001) with Dunnett’s multiple comparisons, *Gal4*/*UAS* versus *Gal4*: \*\*\*\**p*<0.0001, *Gal4*/*UAS* versus *UAS*: \*\*\*\**p*<0.0001. Middle panel: One-way ANOVA (*p*<0.0001) with Dunnett’s multiple comparisons, *Gal4*/*UAS* versus *Gal4*: \*\*\*\**p*<0.0001, *Gal4*/*UAS* versus *UAS*: \*\*\*\**p*<0.0001. Right panel: One-way ANOVA (*p*=0.0588). **A’**. Left panel: One-way ANOVA (*p*<0.0001) with Dunnett’s multiple comparisons, *Gal4*/*UAS* versus *Gal4*: \*\*\*\**p*<0.0001, *Gal4*/*UAS* versus *UAS*: \*\*\*\**p*<0.0001. Middle panel: Brown-Forsythe ANOVA (*p*<0.0001) with Dunnett’s T3 multiple comparisons, *Gal4*/*UAS* versus *Gal4*: \*\*\*\**p*<0.0001, *Gal4*/*UAS* versus *UAS*: \*\*\*\**p*<0.0001. Right panel: Kruskal-Wallis test (p=0.0038) with Dunn’s multiple comparisons, *Gal4*/*UAS* versus *Gal4*: \**p*=0.0141, *Gal4*/*UAS* versus *UAS*: \*\**p*=0.0038. **B**. One-way ANOVA (*p*<0.0001) with Dunnett’s multiple comparisons, *Gal4*/*UAS* versus *Gal4*: \**p*=0.0398, *Gal4*/*UAS* versus *UAS*: \*\*\*\**p*<0.0001. **B’**. One-way ANOVA (*p*<0.0001) with Dunnett’s multiple comparisons, *Gal4*/*UAS* versus *Gal4*: \*\*\*\**p*<0.0001, *Gal4*/*UAS* versus *UAS*: \*\*\*\**p*<0.0001. **C**. One-way ANOVA (*p*=0.0018) with Dunnett’s multiple comparisons, *Gal4*/*UAS* versus *Gal4*: \**p*=0.0209, *Gal4*/*UAS* versus *UAS*: \*\**p*=0.0012. **C’**. Brown-Forsythe ANOVA (*p*=0.0078) with Dunnett’s T3 multiple comparisons, *Gal4*/*UAS* versus *Gal4*: \*\**p*=0.0090, *Gal4*/*UAS* versus *UAS*: *p*=0.1720. **F**. One-way ANOVA (*p*=0.0002) with Dunnett’s multiple comparisons, *Gal4*/*UAS* versus *Gal4*: \*\**p*=0.0038, *Gal4*/*UAS* versus *UAS*: *p*=0.2004. **F’**. One-way ANOVA (*p*<0.0001) with Dunnett’s multiple comparisons, *Gal4*/*UAS* versus *Gal4*: \*\*\*\**p*<0.0001, *Gal4*/*UAS* versus *UAS*: \*\*\*\**p*<0.0001. **G**. One-way ANOVA (*p*=0.0067) with Šídák’s multiple comparisons, no treatment: *p*=0.9716, absorption: *p*=0.9423, metabolism: *p*=0.7767.

Our findings indicated that glutamatergic tolerance neurons are present in both the *R82F12* and the *R18H11 enhancer-Gal4* patterns, and that it is possible that the glutamatergic DN1 circadian neurons are tolerance neurons. Thus, we asked if the *R82F12 enhancer-Gal4* expression pattern includes the DN1 neurons. The DN1p neurons co-express the neuropeptide DH31 and glutamate, and the DN1a neurons co-express the neuropeptide CCHa1 and glutamate (Kunst et al., 2014; Fujiwara et al., 2018). *R82F12* did not express in either the DN1p or the DN1a clusters (**Figure 3D,D’,E,E’**). To further test for possible overlap in *R82F12* and *R18H11* glutamatergic tolerance neurons, we created a *split-Gal4*, with the *R82F12* enhancer driving expression of the GAL4 activation domain (AD) and *R18H11* driving expression of the GAL4 DNA-binding domain (DBD). If glutamatergic tolerance neurons are shared between *R82F12* and *R18H11*, then reconstituted functional GAL4 in the *R82F12*⋂*R18H11 split-Gal4* overlap expressing *vGlut* RNAi should result in decreased tolerance. No decrease in tolerance was observed (**Figure 3F,F’**). Finally, we found that decreased glutamatergic signaling in the DN1 neurons did not affect the absorption or metabolism of ethanol, indicating that glutamate release from DN1 neurons is critical for the behavioral response to ethanol (**Figure 3G**). Thus *R82F12* and *R18H11* contain different glutamatergic tolerance neurons. We chose to focus on the DN1 neurons, because tools exist to precisely separate the the DN1 neuron groups, and because prior research has assigned functions to them.

A panel of three additional *enhancer-Gal4* transgenes was used to determine which, if any, of the DN1 neuron groups promote ethanol tolerance (**Figure 4A**). *tim-Gal4* expresses in all clock neurons, including the DN1a and DN1p neurons, whereas *R16C05-Gal4* expresses specifically in the DN1a neurons and *R51H05-Gal4* express specifically in the DN1p neurons (Kunst et al., 2014; Schubert et al., 2018). RNAi against *vGlut* in *tim-Gal4* and *R16C05-Gal4* reduced tolerance, whereas expression in *R51H05-Gal4* did not (**Figure 4B,B’**). Thus, the pair of DN1a circadian clock neurons promote rapid tolerance development through glutamatergic signaling.

**Figure 4.**
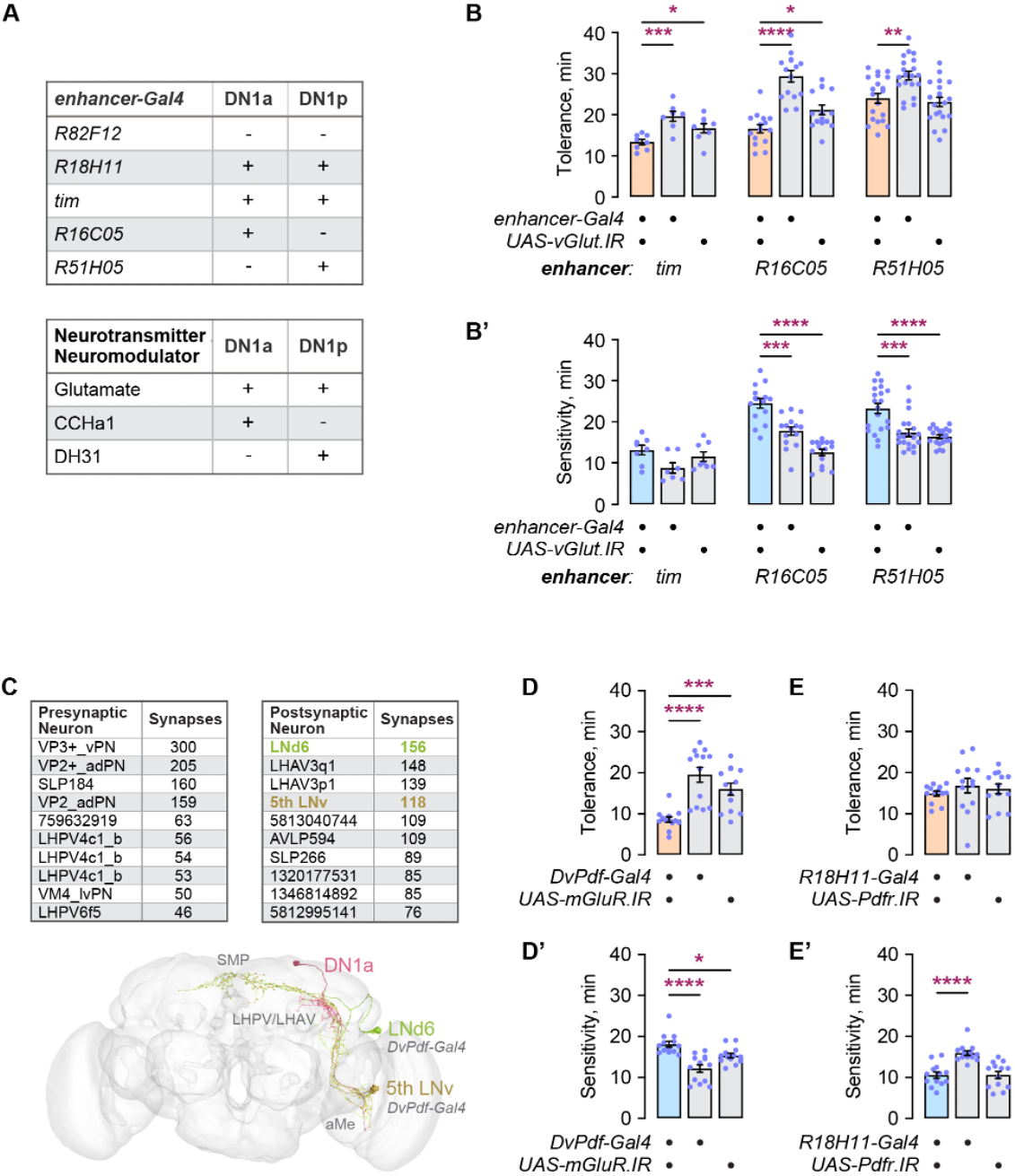
DN1a and downstream evening circadian neurons promote rapid tolerance. **A**. Presence (+) and absence (-) of expression of *enhancer-Gal4*s and neurotransmitter/neuromodulators in the DN1a and DN1p dorsal clock neurons. **B,B’**. Reduction of glutamate release with vesicular glutamate transporter *vGlut* RNAi in *enhancer-Gal4* patterns that include one or both of the dorsal group clock neurons, effects on rapid tolerance (B) and sensitivity (B’). **C**. Ten presynaptic (left) and postsynaptic (right) neurons that make the greatest number of synapses with the DN1a neurons, expressed as the sum total for the right pair of DN1a neurons in the hemibrain electron microscopy reconstruction. The diagram below depicts the morphology of one of the pair of DN1a neurons and the LNd6 and 5th LNv postsynaptic neurons in the full adult female brain (FAFB) electron microscopy reconstruction. SMP: superior medial protocerebrum; LHPV/LHAV: lateral horn posterior/anterior ventral; aMe: accessory medulla. *DvPdf-Gal4* is expressed in the LNd6 and 5th LNv neurons. **D,D’**. Reduction of metabotropic glutamate receptor *mGluR* expression in *DvPdf-Gal4* expressing neurons, effect on rapid tolerance (D) and sensitivity (D’). **E,E**’. Reduction of PDF receptor Pdfr expression in *R18H11-Gal4* expressing neurons, effect on rapid tolerance (E) and sensitivity (E’). *Statistics*: **B**-**B’,D**-**E’**: Quantitative data are mean±SEM. **B**. Left panel: One-way ANOVA (*p*=0.0013) with Dunnett’s multiple comparisons, *Gal4*/*UAS* versus *Gal4*: \*\*\**p*=0.0007, *Gal4*/*UAS* versus *UAS*: \**p*=0.0493. Middle panel: One-way ANOVA (*p*<0.0001) with Dunnett’s multiple comparisons, *Gal4*/*UAS* versus *Gal4*: \*\*\*\**p*<0.0001, *Gal4*/*UAS* versus *UAS*: \**p*=0.0195. Right panel: One-way ANOVA (*p*=0.0002) with Dunnett’s multiple comparisons, *Gal4*/*UAS* versus *Gal4*: \*\**p*=0.0018, *Gal4*/*UAS* versus *UAS*: *p*=0.8003. **B’**. Left panel: One-way ANOVA (*p*=0.0561). Middle panel: One-way ANOVA (*p*<0.0001) with Dunnett’s multiple comparisons, *Gal4*/*UAS* versus *Gal4*: \*\*\*\**p*<0.0001, *Gal4*/*UAS* versus *UAS*: \*\*\*\**p*<0.0001. Right panel: Brown-Forsythe ANOVA (*p*<0.0001) with Dunnett’s T3 multiple comparisons, *Gal4*/*UAS* versus *Gal4*: \*\*\**p*=0.0009, *Gal4*/*UAS* versus *UAS*: \*\*\*\**p*<0.0001. **D**. Brown-Forsythe ANOVA (*p*<0.0001) with Dunnett’s T3 multiple comparisons, *Gal4*/*UAS* versus *Gal4*: \*\*\*\**p*<0.0001, *Gal4*/*UAS* versus *UAS*: \*\*\**p*=0.0005. **D’**. One-way ANOVA (*p*<0.0001) with Dunnett’s multiple comparisons, *Gal4*/*UAS* versus *Gal4*: \*\*\*\**p*<0.0001, *Gal4*/*UAS* versus *UAS*: \**p*=0.0221. **E**. Brown-Forsythe ANOVA (*p*=0.5724). **E’**. One-way ANOVA (*p*<0.0001) with Dunnett’s multiple comparisons, *Gal4*/*UAS* versus *Gal4*: \*\*\*\**p*<0.0001, *Gal4*/*UAS* versus *UAS*: *p*=0.9985.

DN1a neurons set arousal thresholds to brief pulses of light during evening sleep, transmit temperature information into the circadian circuitry, and may promote photoentrainment of the clock by the visual system (Li et al., 2018; Alpert et al., 2020; Song et al., 2021). The synaptic connectivity of the DN1a neurons is well characterized from electron micrograph reconstructions and computational detection of synapses in the adult Drosophila brain (Scheffer et al., 2020; Shafer et al., 2022). Fifty neurons make five or more input synapses with each of the two DN1a neurons per brain hemisphere, and 58-63 neurons are postsynaptic to the DN1a neurons by the same criteria. The top ten presynaptic and postsynaptic neurons are listed in **Figure 4C**. In particular, two circadian neurons that control evening activity, the LNd6 and 5^th^ LNv (previously named the 5^th^ s-LNv) are postsynaptic to the DN1a neurons, providing a connection to circadian pacemakers. The DN1a neurons make synaptic connections to these E2 evening neurons in two regions of the brain, the accessory medulla (aMe) and the Lateral Horn Anterior/Posterior Ventral (LHPV/LHAV) (**Figure 4C**, lower panel). The LNd6 and 5^th^ LNv are similar anatomically and by synaptic connectivity patterns, and they both extend branches into the Superior Medial Protocerebrum (SMP), a higher order processing region of the brain (Shafer et al., 2022).

We performed a test to ask if glutamatergic transmission from the DN1a neurons to the E2 evening neurons might be critical for rapid ethanol tolerance. The single metabotropic glutamate receptor in Drosophila, mGluR, functions in the LNd neurons that includes LNd6 (Guo et al., 2016). We reduced *mGluR* expression in the LNd6 and 5^th^ LNv neurons (as well as other lateral clock neurons) using the *DvPdf-Gal4* driver, and found that tolerance was reduced (**Figure 4D,D’**). The DN1a neurons are reported to be postsynaptic to PDF peptidergic morning clock neurons through the PDF receptor PDFR (Song et al., 2021). Reduction of *Pdfr* expression in the DN1a neurons, however, had no effect on tolerance (**Figure 4E,E’**). Thus, E2 evening circadian neurons are important for the promotion of tolerance development; tolerance information may be transmitted from the glutamatergic DN1a sleep regulatory neurons via the metabotropic glutamate receptor to the E2 neurons.

## Discussion

Our main conclusion is that rapid ethanol tolerance is encoded in part in the two glutamatergic DN1a dorsal clock neurons and the postsynaptic E2 clock neurons LNd6 and 5^th^ LNv. These four clock neurons organize locomotor activity states and aspects of sleep. There exist intriguing relationships between the effects of ethanol, the circadian clock, and sleep in Drosophila and mammals (Spanagel et al., 2005; Perreau-Lenz et al., 2009; Parekh et al., 2015; López-Muciño et al., 2022). Our findings identify specific glutamatergic neurons that likely encode aspects of these relationships.

Glutamatergic transmission in Drosophila was indirectly implicated in tolerance in prior findings. The fly HOMER1/HOMER2 ortholog Homer is reduced in expression by ethanol exposure, and Homer promotes rapid tolerance development (Urizar et al., 2007). Homer proteins function as scaffolding proteins at glutamate receptor postsynaptic densities (Castelli et al., 2017). In mammals, alterations to glutamatergic neurotransmission are critical for ethanol sensitivity and ethanol adaptations, including rapid tolerance (Lindsay et al., 2014; Burnett et al., 2016). Moreover, glutamatergic signaling in mammals is important for aspects of central circadian pacemaker function that is altered by ethanol (Lindsay and Prosser, 2018).

The DN1a dorsal clock neurons regulate aspects of sleep, with their known sleep roles regulated by environmental input. First, cold temperature suppresses DN1a neural activity through activation of cold-temperature-encoding TPN-II second order projection neurons, resulting in a shift to earlier daytime sleep (Alpert et al., 2020, 2022; Chen et al., 2022). This role requires a functioning clock in the DN1a neurons. Second, DN1a neuronal activity specifically promotes nighttime sensitivity to light-induced locomotor startle responses (Song et al., 2021). At night the DN1a neurons increase their arborization and synaptic number in the accessory medulla region, and this remodeling is critical for the circadian time that light can alter startle sensitivity. Acute ethanol exposure alters the expression level of genes encoding presynaptic proteins, and we showed that these genes are critical for rapid tolerance development. Thus, a potential mechanism for tolerance encoding directly in the DN1a neurons is an impact on synaptic plasticity.

The DN1p dorsal clock neurons also regulate sleep, however glutamatergic signaling from this group of neurons is not required for rapid tolerance. Distinct DN1p neurons are sleep promoting and sleep suppressing, and there may exist a parallel segregation for tolerance (Lamaze et al., 2018; Jin et al., 2021). Moreover, despite being glutamatergic, DN1p wake promotion is driven by the neuropeptide CNMa and sleep promotion is driven by another neuropeptide, AstC (Díaz et al., 2018; Jin et al., 2021). Thus, while our current data does not support a potential connection between ethanol tolerance and sleep regulation in the DN1p neurons, more targeted experiments are needed to make a formal conclusion.

The E2 neurons were recently well-segregated as an anatomically, and thus likely functionally, separate group of clock neurons (Schubert et al., 2018; Shafer et al., 2022). Combined with prior findings, it is now clear that the E2 neurons share very similar gene expression profiles. Ion transport peptide (ITP) is expressed in both E2 neurons – the only two clock neurons to express the peptide (Hermann-Luibl et al., 2014). ITP in the clock neurons promotes daytime siestas and nighttime sleep redundantly with the Pigment Dispersing Factor (PDF) neuropeptide, and it separately suppresses nighttime locomotor activity. Thus, both the DN1a neurons and the E2 neurons control aspects of sleep and circadian regulation of locomotor activity levels, strengthening the likelihood that rapid tolerance is tied to one or both behaviors. The E2 neuron outputs include other clock neurons, including the E1 LNds and the DN1ps, but not the DN1as, as well as many non-clock neurons (Shafer et al., 2022). Interestingly, the E2 neurons are not tightly coupled to the M group circadian pacemaker cells that set morning behaviors, supporting the notion that the E2 neurons shape aspects of circadian behavior other than the daily cycle itself (Yao and Shafer, 2014). Hence rapid tolerance encoding may involve a complex clock circuit, or it may exit the circadian clock network via the E2 LNd6 or 5^th^ LNv. The mushroom bodies and the ellipsoid bodies, other sites of rapid ethanol tolerance encoding, also have specific roles in sleep, suggesting that the tie between rapid tolerance and sleep is strongly interrelated (Sitaraman et al., 2015b, 2015a; Aleman et al., 2021).

The Drosophila clock is previously implicated in ethanol sensitivity and rapid ethanol tolerance. Mutation of the key pacemaker genes *per, tim*, and *cyc* completely blocks rapid tolerance, whereas mutations in the *Clk* pacemaker gene did not (Pohl et al., 2013). Hence disruption to the central pacemaker blocks rapid tolerance. Sleep and sleep rebound persist in *per* and *tim* mutants, but sleep is distributed differently over 24 hr (Hendricks et al., 2003).

Sleep and rapid tolerance are genetically and behaviorally connected in flies. Mutation of the learning and memory gene *dunce* results in sleep deficits and a failure to develop rapid tolerance (Ruppert et al., 2017). The anatomical site of action for *dunce* in tolerance is defined by the *NP6510-Gal4* transgene that is not yet characterized for its expression in the clock neurons. Additionally, a subset of histone demethylases of JmjC class regulate both sleep and rapid tolerance (Pinzón et al., 2017; Shalaby et al., 2018). Sleep deprivation for 1 day prior to tolerance induction appears to increase rapid tolerance measured at 4 hr, but it reduces tolerance at 24 hr (De Nobrega et al., 2022). However, rapid tolerance does not appear to be regulated by circadian time (van der Linde and Lyons, 2011; De Nobrega and Lyons, 2016). Thus, current evidence supports a role for ethanol in regulating sleep that is causally connected to rapid tolerance development, however the clock does not appear to reciprocally regulate rapid tolerance development.

Finally, ethanol sensitivity appears to map broadly to anatomical sites in the Drosophila brain, but in a different manner as compared to rapid tolerance. For example, glutamatergic promotion of ethanol sensitivity appears to map to neurons that are shared between *R18H11-Gal4* and *R82F12-Gal4*, whereas the rapid tolerance neurons in these two *enhancer-Gal4* patterns mapped to distinct groups of neurons. Additionally, glutamatergic ethanol sensitivity appeared to map to non-clock neurons, since *tim-Gal4* reduction in *vGlut* expression did not affect ethanol sensitivity. These findings are in accordance with prior studies indicating functional separation of the ethanol sensitivity and rapid tolerance neural circuitry (Chvilicek et al., 2020).

## Methods

### Drosophila culturing and strains

Drosophila melanogaster strains were reared on food composed of molasses (9%), cornmeal (6.75%), yeast (1.7%), and agar (1.2%) food at 25°C and 60% humidity on a 16:8 hour light/dark cycle (Darwin Chambers, MO). All strains were outcrossed for at least five generations to the Berlin genetic background strain carrying the *w*^*1118*^ marker mutation. The RNAi transgenes used in this study were validated previously (Guo et al., 2016; Aguilar et al., 2017; Nandi et al., 2017; Newman et al., 2022). Strains are listed in **Table S1**.

### Behavioral studies

Parental crosses were set up in Drosophila culturing bottles containing 50 mL of food. After two days the parents were removed. Fourteen days later 0-3 day old genetically identical adult male progeny were collected in groups of 20 (n=1) and allowed to recover from CO2 anesthesia for 2 days. Ethanol sensitivity and rapid tolerance were measured as previously described (Engel et al., 2016). Briefly, groups were exposed to 55% ethanol vapor or 100% humidified air. 55% ethanol is an intermediate ethanol dose that results in submaximal rapid tolerance and in 50% sedation in 12-20 min (Kong et al., 2010a). The number of flies that lost the righting reflex were counted at 10 min intervals. The time to 50% sedation (ST50) was calculated for each group (E1). Flies were allowed to rest for 3.5 hr and re-exposed to an identical concentration of ethanol vapor (E2). Rapid tolerance was calculated as the difference in ST50, E2-E1.

### Ethanol absorption and metabolism

Groups of 20 flies were exposed to 30 min of 20% ethanol to avoid sedation, then frozen in liquid nitrogen either immediately for “absorption” samples, or 30 min later for “metabolism” samples. After homogenization in 50 mM Tris-HCl, pH 7.5, ethanol concentrations were measured using the NAD-ADH Reagent kit following the manufacturer’s protocol (Sigma-Aldrich, N7160). The 340 nm spectrophotometric absorbance values were converted to mM and adjusted by the estimated 1 μL volume of an average fly to calculate ethanol concentrations.

### Immunohistochemistry

Adult fly brains were dissected in PBS with 0.05% Triton X-100 (PBT), fixed overnight at 4°C in PBT with 2% paraformaldehyde, blocked in PBS with 0.5% Triton X-100 with 5% normal goat serum and 0.5% bovine serum albumin (HDB), and immunostained as described previously (Kong et al., 2010b). Antibodies and their concentrations are listed in **Table S1**. The brains were mounted in Vectashield (Vector Laboratories) and imaged on a Zeiss LSM-880 confocal microscope. Image stacks were processed in Fiji, and brightness and contrast were adjusted in Photoshop CC 2022 (Adobe).

### Statistical analysis

Experimental and genetically-matched or treatment-matched controls were tested in the same session in a balanced experimental design. Experiments were repeated across days with progeny from repeat parental crosses, and data from all days and crosses were collated together without between-day adjustments. Untransformed (raw) data was used for statistical analysis. Where the experimental group was compared to two or more control groups, significance was only interpreted when all controls were different from the experimental. GraphPad Prism 9.5.0 was used for one-way ANOVA with Tukey’s post hoc test for normally distributed data, Kruskal-Wallis test with Dunn’s post hoc test for nonparametric data, and Brown-Forsythe test with Dunnett’s post hoc for data that fails the Shapiro-Wilk normality test. Significance indicators on the figures indicate the results of post hoc tests for significant effects by ANOVA (**** for *p*≤0.0001; *** for *p*≤0.001; ** for *p*≤0.01; * for *p*≤0.05; and ns for *p*>0.05). Error bars represent the standard error of the mean (SEM).

## Acknowledgments

We thank members of the Wolf lab for intellectual and editing support. This work was supported by R21 AA029178 and R21/R33 AA028352 (F.W.W).

---

**Table S1.**
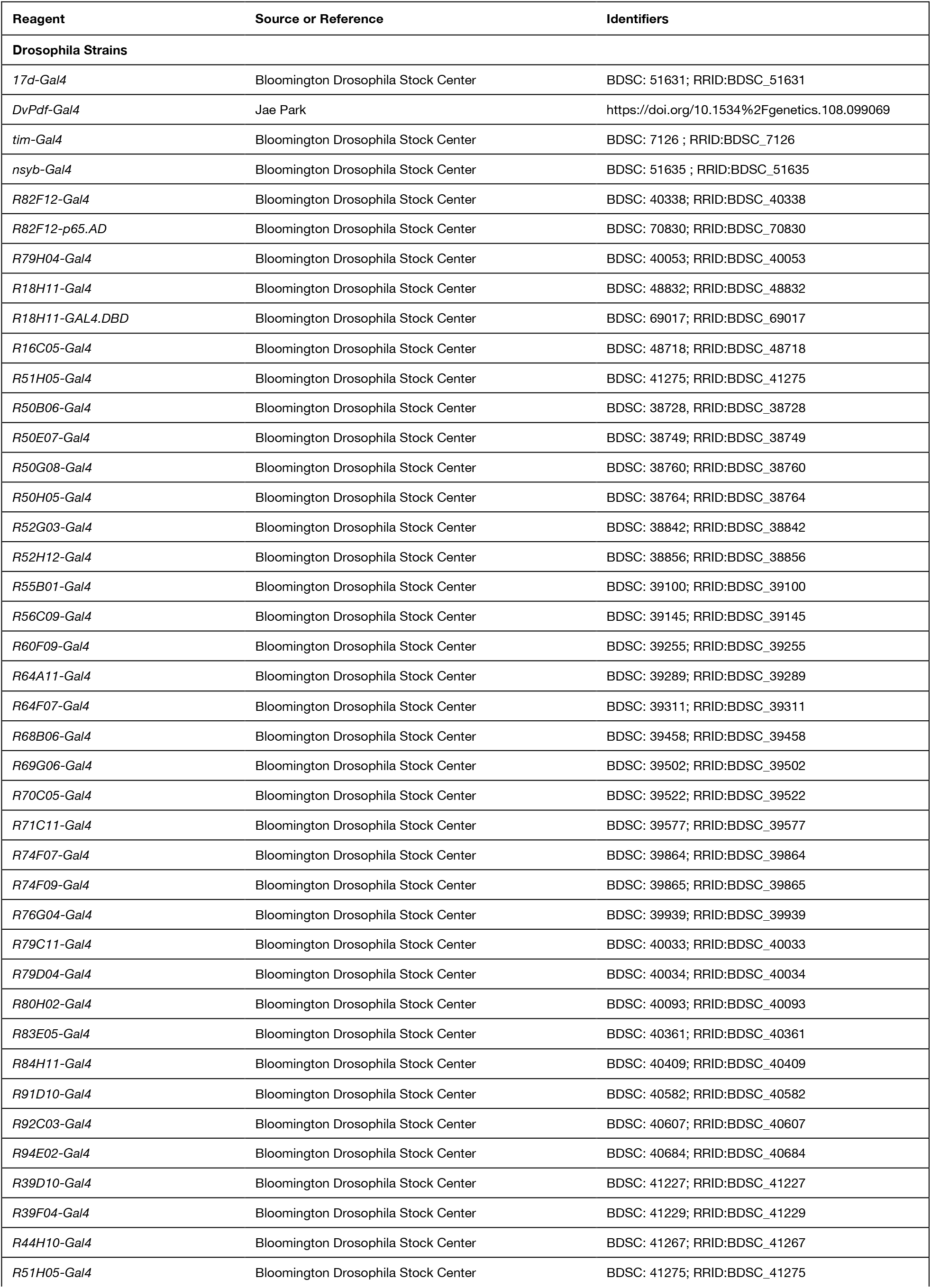
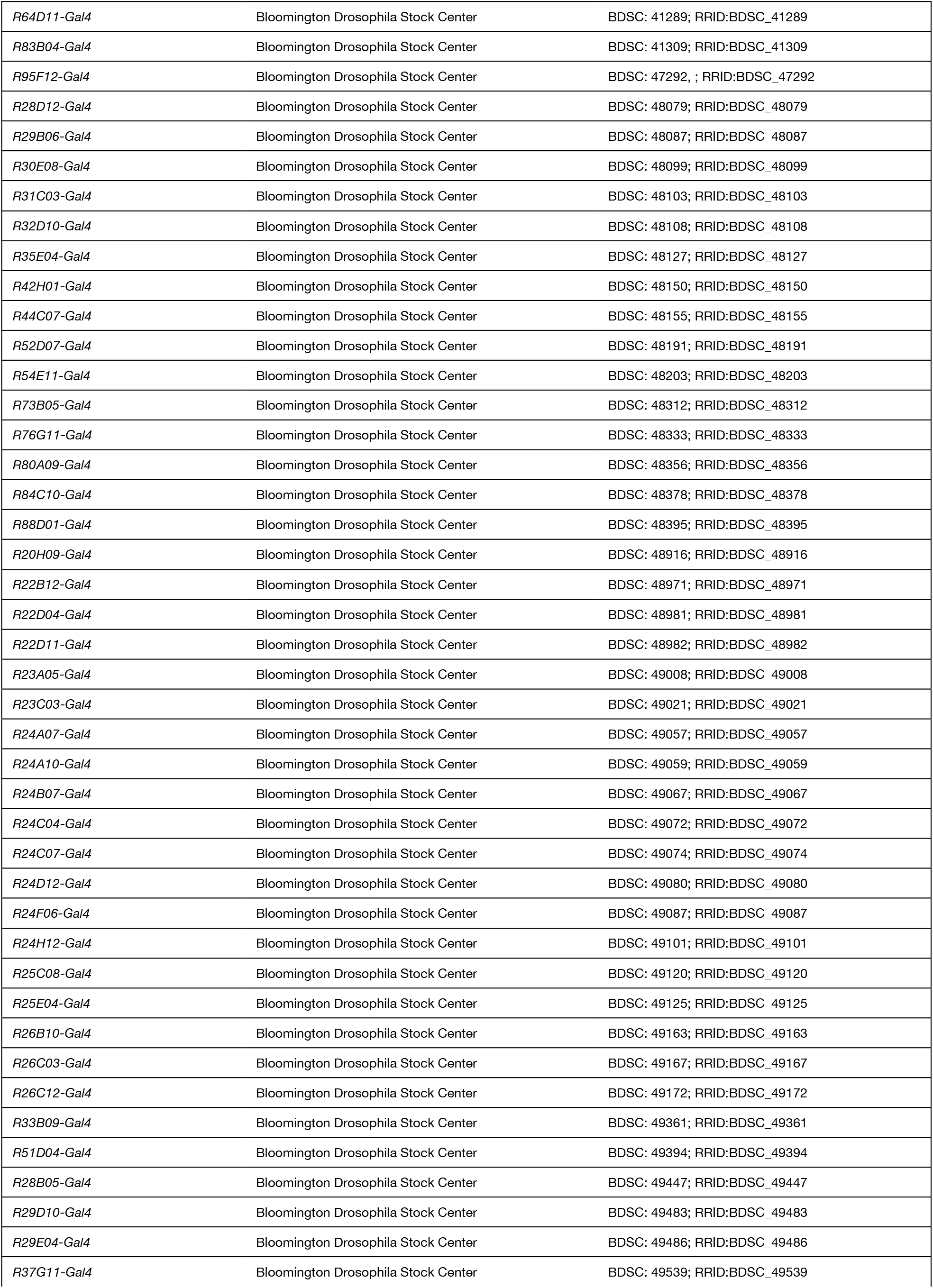
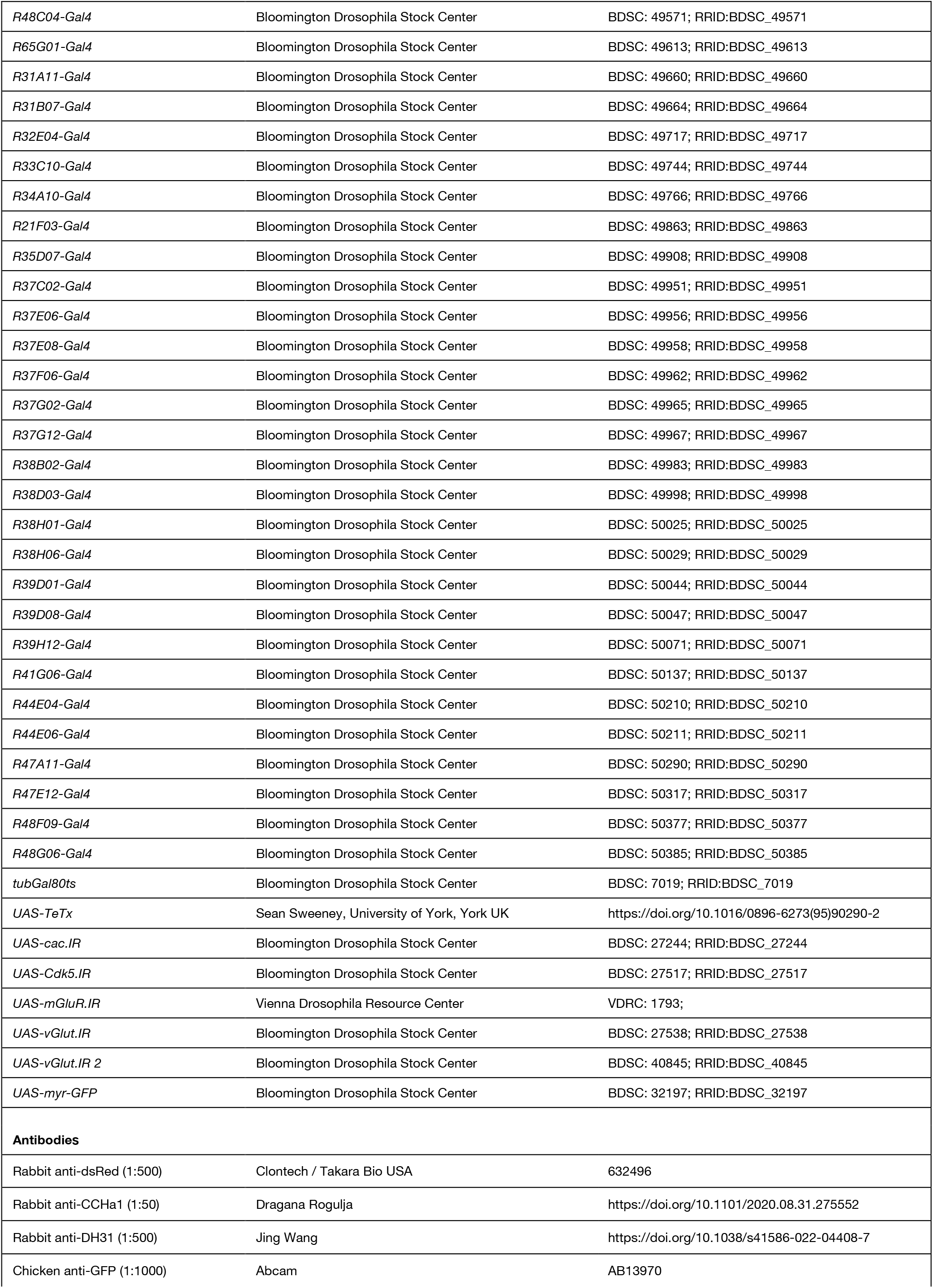
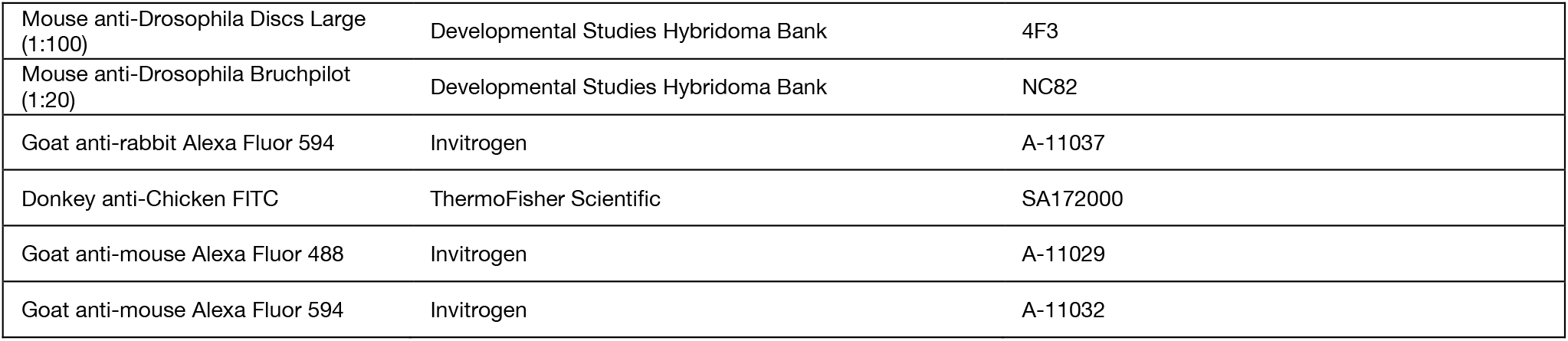
Reagents.

## References

Aguilar JI et al. (2017) Neuronal Depolarization Drives Increased Dopamine Synaptic Vesicle Loading via VGLUT. Neuron 95:1074-1088.e7.

Aleman A, Omoto JJ, Singh P, Nguyen B-C, Kandimalla P, Hartenstein V, Donlea JM (2021) Opposing subclasses of Drosophila ellipsoid body neurons promote and suppress sleep. :2021.10.19.464469 Available at: https://www.biorxiv.org/content/10.1101/2021.10.19.464469v1 [Accessed January 14, 2023].

Alpert MH, Frank DD, Kaspi E, Flourakis M, Zaharieva EE, Allada R, Para A, Gallio M (2020) A Circuit Encoding Absolute Cold Temperature in Drosophila. Curr Biol 30:2275-2288.e5.

Alpert MH, Gil H, Para A, Gallio M (2022) A thermometer circuit for hot temperature adjusts Drosophila behavior to persistent heat. Curr Biol 32:4079-4087.e4.

Andreani T, Rosensweig C, Sisobhan S, Ogunlana E, Kath W, Allada R (2022) Circadian programming of the ellipsoid body sleep homeostat in Drosophila. Elife 11:e74327.

Berger KH, Heberlein U, Moore MS (2004) Rapid and chronic: two distinct forms of ethanol tolerance in Drosophila. Alcohol Clin Exp Res 28:1469–1480.

Berkel TDM, Pandey SC (2017) Emerging Role of Epigenetic Mechanisms in Alcohol Addiction. Alcohol Clin Exp Res 41:666–680.

Burnett EJ, Chandler LJ, Trantham-Davidson H (2016) Glutamatergic plasticity and alcohol dependence-induced alterations in reward, affect and cognition. Prog Neuropsychopharmacol Biol Psychiatry 65:309–320.

Castelli V, Brancato A, Cavallaro A, Lavanco G, Cannizzaro C (2017) Homer2 and Alcohol: A Mutual Interaction. Front Psychiatry 8:268.

Chen S-C, Tang X, Goda T, Umezaki Y, Riley AC, Sekiguchi M, Yoshii T, Hamada FN (2022) Dorsal clock networks drive temperature preference rhythms in Drosophila. Cell Rep 39:110668.

Chvilicek MM, Titos I, Rothenfluh A (2020) The Neurotransmitters Involved in Drosophila Alcohol-Induced Behaviors. Front Behav Neurosci 14:607700.

Cowmeadow RB, Krishnan HR, Atkinson NS (2005) The slowpoke gene is necessary for rapid ethanol tolerance in Drosophila. Alcohol Clin Exp Res 29:1777–1786.

De Nobrega AK, Lyons LC (2016) Circadian Modulation of Alcohol-Induced Sedation and Recovery in Male and Female Drosophila. J Biol Rhythms 31:142–160.

De Nobrega AK, Noakes EJ, Storch NA, Mellers AP, Lyons LC (2022) Sleep Modulates Alcohol Toxicity in Drosophila. Int J Mol Sci 23:12091.

Díaz MM, Schlichting M, Abruzzi KC, Long X, Rosbash M (2018) Allatostatin-C/AstC-R2 Is a Novel Pathway to Modulate the Circadian Activity Pattern in Drosophila. Curr Biol.

Engel GL, Marella S, Kaun KR, Wu J, Adhikari P, Kong EC, Wolf FW (2016) Sir2/Sirt1 Links Acute Inebriation to Presynaptic Changes and the Development of Alcohol Tolerance, Preference, and Reward. J Neurosci 36:5241–5251.

Fadda F, Rossetti ZL (1998) Chronic ethanol consumption: From neuroadaptation to neurodegeneration. Progress in Neurobiology 56:385–431.

Fujiwara Y, Hermann-Luibl C, Katsura M, Sekiguchi M, Ida T, Helfrich-Förster C, Yoshii T (2018) The CCHamide1 Neuropeptide Expressed in the Anterior Dorsal Neuron 1 Conveys a Circadian Signal to the Ventral Lateral Neurons in Drosophila melanogaster. Front Physiol 9:1276.

Galili DS, Jefferis GS, Costa M (2022) Connectomics and the neural basis of behaviour. Curr Opin Insect Sci 54:100968.

Ghezzi A, Al-Hasan YM, Krishnan HR, Wang Y, Atkinson NS (2013a) Functional mapping of the neuronal substrates for drug tolerance in Drosophila. Behav Genet 43:227–240.

Ghezzi A, Krishnan HR, Lew L, Prado FJ, Ong DS, Atkinson NS (2013b) Alcohol-induced histone acetylation reveals a gene network involved in alcohol tolerance. PLoS Genet 9:e1003986.

Godenschwege TA et al. (2004) Flies lacking all synapsins are unexpectedly healthy but are impaired in complex behaviour. Eur J Neurosci 20:611–622.

Guo F, Yu J, Jung HJ, Abruzzi KC, Luo W, Griffith LC, Rosbash M (2016) Circadian neuron feedback controls the Drosophila sleep--activity profile. Nature 536:292–297.

Hendricks JC, Lu S, Kume K, Yin JCP, Yang Z, Sehgal A (2003) Gender dimorphism in the role of cycle (BMAL1) in rest, rest regulation, and longevity in Drosophila melanogaster. J Biol Rhythms 18:12–25.

Hermann-Luibl C, Yoshii T, Senthilan PR, Dircksen H, Helfrich-Förster C (2014) The ion transport peptide is a new functional clock neuropeptide in the fruit fly Drosophila melanogaster. J Neurosci 34:9522–9536.

Jin X, Tian Y, Zhang ZC, Gu P, Liu C, Han J (2021) A subset of DN1p neurons integrates thermosensory inputs to promote wakefulness via CNMa signaling. Curr Biol 31:2075-2087.e6.

Kang YY, Wachi Y, Engdorf E, Fumagalli E, Wang Y, Myers J, Massey S, Greiss A, Xu S, Roman G (2020) Normal Ethanol Sensitivity and Rapid Tolerance Require the G Protein Receptor Kinase 2 in Ellipsoid Body Neurons in Drosophila. Alcohol Clin Exp Res 44:1686–1699.

Kaun KR, Azanchi R, Maung Z, Hirsh J, Heberlein U (2011) A Drosophila model for alcohol reward. Nat Neurosci 14:612–619.

Kong EC, Allouche L, Chapot PA, Vranizan K, Moore MS, Heberlein U, Wolf FW (2010a) Ethanol-regulated genes that contribute to ethanol sensitivity and rapid tolerance in Drosophila. Alcohol Clin Exp Res 34:302–316.

Kong EC, Woo K, Li H, Lebestky T, Mayer N, Sniffen MR, Heberlein U, Bainton RJ, Hirsh J, Wolf FW (2010b) A pair of dopamine neurons target the D1-like dopamine receptor DopR in the central complex to promote ethanol-stimulated locomotion in Drosophila. PLoS One 5:e9954.

Kunst M, Hughes ME, Raccuglia D, Felix M, Li M, Barnett G, Duah J, Nitabach MN (2014) Calcitonin gene-related peptide neurons mediate sleep-specific circadian output in Drosophila. Curr Biol 24:2652–2664.

Lamaze A, Krätschmer P, Chen K-F, Lowe S, Jepson JEC (2018) A Wake-Promoting Circadian Output Circuit in Drosophila. Curr Biol.

Li M-T, Cao L-H, Xiao N, Tang M, Deng B, Yang T, Yoshii T, Luo D-G (2018) Hub-organized parallel circuits of central circadian pacemaker neurons for visual photoentrainment in Drosophila. Nat Commun 9:4247.

Lindsay JH, Glass JD, Amicarelli M, Prosser RA (2014) The mammalian circadian clock in the suprachiasmatic nucleus exhibits rapid tolerance to ethanol in vivo and in vitro. Alcohol Clin Exp Res 38:760–769.

Lindsay JH, Prosser RA (2018) The Mammalian Circadian Clock Exhibits Chronic Ethanol Tolerance and Withdrawal-Induced Glutamate Hypersensitivity, Accompanied by Changes in Glutamate and TrkB Receptor Proteins. Alcohol Clin Exp Res 42:315–328.

López-Muciño LA, García-García F, Cueto-Escobedo J, Acosta-Hernández M, Venebra-Muñoz A, Rodríguez-Alba JC (2022) Sleep loss and addiction. Neurosci Biobehav Rev 141:104832.

Luan H, Diao F, Scott RL, White BH (2020) The Drosophila Split Gal4 System for Neural Circuit Mapping. Front Neural Circuits 14:603397.

McGuire SE, L. PT, Osborn AJ, Matsumoto K, Davis RL (2003) Spatiotemporal rescue of memory dysfunction in Drosophila. Science 302:1765–1768.

Modi MN, Shuai Y, Turner GC (2020) The Drosophila Mushroom Body: From Architecture to Algorithm in a Learning Circuit. Annu Rev Neurosci 43:465–484.

Morozova TV, Anholt RR, Mackay TF (2006) Transcriptional response to alcohol exposure in Drosophila melanogaster. Genome Biol 7:R95.

Nandi N, Tyra LK, Stenesen D, Krämer H (2017) Stress-induced Cdk5 activity enhances cytoprotective basal autophagy in Drosophila melanogaster by phosphorylating acinus at serine437. Elife 6:e30760.

Newman ZL, Bakshinskaya D, Schultz R, Kenny SJ, Moon S, Aghi K, Stanley C, Marnani N, Li R, Bleier J, Xu K, Isacoff EY (2022) Determinants of synapse diversity revealed by super-resolution quantal transmission and active zone imaging. Nat Commun 13:229.

Parekh PK, Ozburn AR, McClung CA (2015) Circadian clock genes: effects on dopamine, reward and addiction. Alcohol 49:341–349.

Park A, Ghezzi A, Wijesekera TP, Atkinson NS (2017) Genetics and genomics of alcohol responses in Drosophila. Neuropharmacology 122:22–35.

Parkhurst SJ, Adhikari P, Navarrete JS, Legendre A, Manansala M, Wolf FW (2018) Perineurial Barrier Glia Physically Respond to Alcohol in an Akap200-Dependent Manner to Promote Tolerance. Cell Rep 22:1647–1656.

Perreau-Lenz S, Zghoul T, de Fonseca FR, Spanagel R, Bilbao A (2009) Circadian regulation of central ethanol sensitivity by the mPer2 gene. Addict Biol 14:253–259.

Peru Y Colón de Portugal RL, Ojelade SA, Penninti PS, Dove RJ, Nye MJ, Acevedo SF, Lopez A, Rodan AR, Rothenfluh A (2014) Long-lasting, experience-dependent alcohol preference in Drosophila. Addict Biol 19:392–401.

Pfeiffer BD, Jenett A, Hammonds AS, Ngo T-TB, Misra S, Murphy C, Scully A, Carlson JW, Wan KH, Laverty TR, Mungall C, Svirskas R, Kadonaga JT, Doe CQ, Eisen MB, Celniker SE, Rubin GM (2008) Tools for neuroanatomy and neurogenetics in Drosophila. Proc Natl Acad Sci USA 105:9715–9720.

Pinzón JH, Reed AR, Shalaby NA, Buszczak M, Rodan AR, Rothenfluh A (2017) Alcohol-Induced Behaviors Require a Subset of Drosophila JmjC-Domain Histone Demethylases in the Nervous System. Alcohol Clin Exp Res 41:2015–2024.

Pohl JB, Ghezzi A, Lew LK, Robles RB, Cormack L, Atkinson NS (2013) Circadian genes differentially affect tolerance to ethanol in Drosophila. Alcohol Clin Exp Res 37:1862–1871.

Ranson DC, Ayoub SS, Corcoran O, Casalotti SO (2020) Pharmacological targeting of the GABAB receptor alters Drosophila’s behavioural responses to alcohol. Addict Biol 25:e12725.

Ruppert M, Franz M, Saratsis A, Velo Escarcena L, Hendrich O, Gooi LM, Schwenkert I, Klebes A, Scholz H (2017) Hangover Links Nuclear RNA Signaling to cAMP Regulation via the Phosphodiesterase 4d Ortholog dunce. Cell Rep 18:533–544.

Sakharkar AJ, Zhang H, Tang L, Shi G, Pandey SC (2012) Histone deacetylases (HDAC)-induced histone modifications in the amygdala: a role in rapid tolerance to the anxiolytic effects of ethanol. Alcohol Clin Exp Res 36:61–71.

Scheffer LK et al. (2020) A connectome and analysis of the adult Drosophila central brain. Elife 9:e57443.

Scholz H, Ramond J, Singh CM, Heberlein U (2000) Functional Ethanol Tolerance in Drosophila. Neuron 28:261–271.

Schubert FK, Hagedorn N, Yoshii T, Helfrich-Förster C, Rieger D (2018) Neuroanatomical details of the lateral neurons of Drosophila melanogaster support their functional role in the circadian system. J Comp Neurol 526:1209–1231.

Seelig JD, Jayaraman V (2015) Neural dynamics for landmark orientation and angular path integration. Nature 521:186–191.

Shafer OT, Gutierrez GJ, Li K, Mildenhall A, Spira D, Marty J, Lazar AA, Fernandez M de la P (2022) Connectomic analysis of the Drosophila lateral neuron clock cells reveals the synaptic basis of functional pacemaker classes. Elife 11:e79139.

Shafer OT, Helfrich-Förster C, Renn SCP, Taghert PH (2006) Reevaluation of Drosophila melanogaster’s neuronal circadian pacemakers reveals new neuronal classes. J Comp Neurol 498:180–193.

Shalaby NA, Pinzon JH, Narayanan AS, Jin EJ, Ritz MP, Dove RJ, Wolfenberg H, Rodan AR, Buszczak M, Rothenfluh A (2018) JmjC domain proteins modulate circadian behaviors and sleep in Drosophila. Sci Rep 8:815.

Sitaraman D, Aso Y, Jin X, Chen N, Felix M, Rubin GM, Nitabach MN (2015a) Propagation of Homeostatic Sleep Signals by Segregated Synaptic Microcircuits of the Drosophila Mushroom Body. Curr Biol 25:2915–2927.

Sitaraman D, Aso Y, Rubin GM, Nitabach MN (2015b) Control of Sleep by Dopaminergic Inputs to the Drosophila Mushroom Body. Front Neural Circuits 9:73.

Song BJ, Sharp SJ, Rogulja D (2021) Daily rewiring of a neural circuit generates a predictive model of environmental light. Sci Adv 7:eabe4284.

Spanagel R, Pendyala G, Abarca C, Zghoul T, Sanchis-Segura C, Magnone MC, Lascorz J, Depner M, Holzberg D, Soyka M, Schreiber S, Matsuda F, Lathrop M, Schumann G, Albrecht U (2005) The clock gene Per2 influences the glutamatergic system and modulates alcohol consumption. Nat Med 11:35–42.

Urizar NL, Yang Z, Edenberg HJ, Davis RL (2007) Drosophila homer is required in a small set of neurons including the ellipsoid body for normal ethanol sensitivity and tolerance. J Neurosci 27:4541–4551.

van der Linde K, Lyons LC (2011) Circadian modulation of acute alcohol sensitivity but not acute tolerance in Drosophila. Chronobiol Int 28:397–406.

Xu S, Chan T, Shah V, Zhang S, Pletcher SD, Roman G (2012) The propensity for consuming ethanol in Drosophila requires rutabaga adenylyl cyclase expression within mushroom body neurons. Genes Brain Behav 11:727–739.

Yao Z, Shafer OT (2014) The Drosophila circadian clock is a variably coupled network of multiple peptidergic units. Science 343:1516–1520.

